# Identification of 55,000 Replicated DNA Methylation QTL

**DOI:** 10.1101/166710

**Authors:** Allan F McRae, Riccardo E Marioni, Sonia Shah, Jian Yang, Joseph E. Powell, Sarah E Harris, Jude Gibson, Anjali K Henders, Lisa Bowdler, Jodie N. Painter, Lee Murphy, Nicholas G Martin, John M Starr, Naomi R Wray, Ian J Deary, Peter M Visscher, Grant W Montgomery

## Abstract

DNA methylation plays an important role in the regulation of transcription. Genetic control of DNA methylation is a potential candidate for explaining the many identified SNP associations with disease that are not found in coding regions. We replicated 52,916 *cis* and 2,025 *trans* DNA methylation quantitative trait loci (mQTL) using methylation measured on Illumina HumanMethylation450 arrays in the Brisbane Systems Genetics Study (n=614 from 177 families) and the Lothian Birth Cohorts of 1921 and 1936 (combined n = 1366). The *trans* mQTL SNPs were found to be over-represented in 1Mbp subtelomeric regions, and on chromosomes 16 and 19. There was a significant increase in *trans* mQTL DNA methylation sites in upstream and 5’ UTR regions. No association was observed between either the SNPs or DNA methylation sites of *trans* mQTL and telomere length. The genetic heritability of a number of complex traits and diseases was partitioned into components due to mQTL and the remainder of the genome. Significant enrichment was observed for height (p = 2.1x10^−10^), ulcerative colitis (p = 2x10^−5^), Crohn’s disease (p = 6x10^−8^) and coronary artery disease (p = 5.5x10^−6^) when compared to a random sample of SNPs with matched minor allele frequency, although this enrichment is explained by the genomic location of the mQTL SNPs.

## INTRODUCTION

DNA methylation plays an important role in transcriptional regulation and is increasingly recognised as having a role in health and disease ^1,2^. The contribution of genetic variation to the inheritance of DNA methylation levels across a range of tissues has been widely demonstrated both through studies investigating the heritability of DNA methylation using twin pairs and families ^3–6^, and through the identification of methylation quantitative trait loci or mQTL acting in both *cis* and *trans* ^7–19^.

As the majority of single nucleotide polymorphisms (SNPs) associated with complex traits and disease are found in non-coding regions ^20^, it is hypothesised that the SNPs act through the perturbation of the regulation of gene-expression. DNA methylation QTL have been associated with other genomic marks that affect gene regulation, including DNase I accessibility and histone modifications ^16,17^, as well as directly with gene-expression ^15,16^, Therefore, they are potential causal variants for disease. Indeed, the overlap between mQTL and disease SNPs has been investigated previously, finding inflation for the number of mQTL in bipolar risk SNPs ^11^, schizophrenia ^18^ and autoimmune disease ^17^.

These published studies indicate that mQTL have an influence in disease risk, however some aspects of the methodological approach in determining the significance of the overlap may be sub-optimal. For example, most identified mQTL have been found using Illumina HumanMethylation arrays, but the analytical methods have not recognised that the measures of DNA methylation are distrubuted non-randomly throughout the genome. Most of the DNA methylation probes on these arrays are located in genic regions, and, given that the majority of mQTL are found in *cis* to DNA methylation sites, the mQTL SNPs are also preferentially located in genic regions. Genic regions are also known to explain a larger proportion of the genetic variation underlying complex traits and disease ^21^. Therefore, any analysis looking into the overlap of mQTL with SNPs identified in genome-wide association studies (GWAS) needs to account for the proportion of methylation sites assessed in different genomic regions. In addition, determining the overlap between a mQTL and disease SNP often uses criteria such as an arbitrary linkage disequilibrium (LD) threshold of r^2^ > 0.8 between the best disease GWAS SNP and the mQTL SNP. This implicitly assumes that a common causal variant for the mQTL and disease is being tagged by two different SNPs, rather than there being two different causal variants.

Here we use two large genomic studies - the Brisbane Systems Genetics Study (BSGS) ^6,22^ and the Lothian Birth Cohorts of 1921 and 1936 (LBC) ^23–25^ - to identify >50,000 mQTL that are replicated at a stringent significance level. These mQTL are then used to partition the genetic variation for complex traits and diseases into components due to mQTL SNPs and the remainder of the genome using LD Score regression ^26,27^ on the summary statistics from large GWAS meta-analyses. This avoids selecting an arbitrary linkage disequilibrium threshold above which mQTL and disease SNPs are considered as overlapping. These analyses are compared to null distributions generated by selecting random sets of SNPs that have been matched by allele frequency or by both allele frequency and genomic annotation.

## RESULTS

### Identification of mQTL

Due to prior evidence showing large *cis* SNP effects on DNA methylation, we firstly tested for association in a window spanning 2Mbp either side of the target CpG site. This window is larger than what is usually considered for *cis* mQTL, but our prior observation of significant *cis* mQTL effects spanning this far in the MHC region on chromosome 6 indicated a larger window is warranted ^6^. This was further justified by noting that the number of *cis* mQTL rapidly drops off to a constant background level between 1 and 2Mbps from the target CpG site (Figure S1).

A total of 62,257 and 61,180 *cis* mQTL were identified in the BSGS and LBC cohorts respectively at a significance threshold of *p* < 10^−11^. While only the most significant SNP for each DNA methylation probe is considered, many of the mQTL are non-independent due to both correlations between DNA methylation levels for probes separated by small distances and through linkage disequilibrium between SNPs. Of these, 52,916 (~85%) replicated in the other cohort at Bonferonni corrected significance threshold of *p* < 10^−6^ and also had SNP effects on DNA methylation in the same direction in the other cohort. The correlation of *cis* mQTL effect sizes between the two cohorts was 0.97. Thus we have stringently replicated *cis* mQTL for more than 13% of the methylation sites tested.

*Trans* mQTL were defined using a more stringent significance threshold of *p* < 10^−13^ to account for the extra multiple testing burden from testing association with the whole genome. The number of significant *trans* mQTL found in the BSGS and LBC was 2,454 and 2,048 respectively. Of these, 2,025 replicated in the other cohort with a Bonferonni corrected p-value of *p* < 10^−5^ and also had the same direction of effect. The correlation in *trans* mQTL effect sizes across the two cohorts was 0.91. The location of the replicated mQTL are given in Figure 1. The extremely high replication rate for both *cis*- and *trans*-mQTL in independent samples demonstrates the high quality of the data and reliability of the results.

**Figure 1:** Location of replicated mQTL across the genome. Each point represents a replicated mQTL with the position of the CpG site on the X-axis and the SNP location on the Y-axis. Chromosome boundaries are indicated with dashed lines. The diagonal line shows an abundance of *cis* mQTL throughout the genome. Also visible are horizontal bands of *trans* mQTL in the telomeric regions of the chromosomes. See also Figure S1.

The proportion of phenotypic variation in DNA methylation levels explained by all replicated mQTL in the LBC cohort is given in Figure 2. As expected from QTL identified using limited sample sizes (as compared to contemporary GWAS for complex traits and disease), the phenotypic variation explained by the mQTL is very large, with 8% of *cis* mQTL explaining greater than 50% of phenotypic variation. While *trans* mQTL still explain a substantial proportion of the phenotypic variance, the overall distribution has fewer mQTL explaining very large amounts of variance. The effect of the “winner’s curse”, where the variance explained by the top SNPs identified in a GWAS is biased upwards, is likely to be small in this study given the stringency of testing and the high replication rate.

**Figure 2:** Proportion of phenotypic variation of DNA methylation levels explained by mQTL in the LBC cohort.

There is potential for SNPs located within DNA methylation probe binding regions to have an effect on the measurement of methylation levels, and thus potentially create false positive mQTL. To address this, we used the 1000Genomes (v3) European samples to identify any genetic variation within a probe site and identified a SNP in 27% of the probes passing QC. It is of note that many of the SNPs identified within probe sequences are rare and would not be in strong linkage disequilibrium with the common (>1% frequency) SNPs used for the GWAS. For *trans* mQTL, it is very unlikely that a SNP in the probe site was associated with the mQTL SNP, particularly given the very stringent significance thresholds that were used for mQTL mapping. This is reflected in 499 (25%) *trans* mQTL having a SNP in the probe site, which is the same as the null proportion of probes that do not have an associated mQTL that have SNPs in their binding site (85,621/342,967). SNPs were found within the probe binding site for 22,267 (42%) of *cis* mQTL. Thus, we can potentially attribute 15% (42% - 27%) of *cis* mQTL to genetic variation within the probe location causing genotype specific measurement error. However, it can also be argued that the majority of *cis* mQTL are found within a very small distance of the probe location, and it would not be surprising for genetic variation very close to a CpG site to have a genuine effect on methylation levels. To take an extreme example, a SNP falling within a CpG site completely disrupts DNA methylation at this site, which occurs for 6,160 (12%) of *cis* mQTL. For this reason, we include all mQTL – regardless of the identification of SNP within the probe site – in the further analyses.

### Genomic Distribution of *Trans* mQTL

From Figure 1, we have an indication that the distribution of *trans* mQTL SNPs is non-randomly located throughout the genome. This is investigated in Figure 3a, which shows there is a large number of *trans* mQTL SNP located on chromosomes 16 and 19 given their respective sizes. This may not be surprising under a polygenic model of inheritance given those chromosomes have a higher gene density than other chromosomes. However, this inflation is beyond that expected given the gene count on those two chromosomes (Figure 3b). The rest of the genome shows a strong correlation between number of genes on a chromosome and the number of *trans* mQTL SNPs, except for chromosome 1 which has fewer *trans* mQTL SNP than expected. Of interest, chromosome 19 contains DNMT1 (DNA methyltransferase 1) that has a role in the establishment and regulation of DNA methylation. Interestingly however, there is no clustering of *trans* mQTL SNPs around its location.

**Figure 3:** Genomic location of *trans* mQTL. (a) a circos plot showing *trans* mQTL occurring throughout the genome. Chromosomes 16 and 19 have a large number of *trans* mQTL SNPs, and this inflation is beyond that expected due to the increased gene density on those chromosomes (b).

There are clear horizontal bands of SNPs in Figure 1, located in the subtelomeric regions of the genome. Indeed, 17.9% of all *trans* mQTL SNP are located in telomeric regions covering the 1Mbp at the end of chromosomes, which represents 1.53% of the genome. There is also some inflation of the numbers of *trans* mQTL methylation probes found in the 1Mbp subtelomeric region (7.0%), but this is primarily due to the increased number of array probes in the subtelomeric region (5.5%) and this inflation is reflected in the number of *cis* mQTL methylation probes also (7.5%). Given the association with *trans* mQTL SNP in telomeric regions, we tested whether the *trans* CpG probes or SNPs were significantly associated with telomere length in the LBC1936 cohort. This identified no inflation of test statistics for either the SNPs or methylation compared to the whole genome (Figure S3).

Unlike *trans* mQTL SNPs, the CpG probe locations showed no clustering across the genome. To investigate a functional role of the *trans* mQTL methylation sites, we annotated the genomic locations of all the array probes tested (Table 1). As expected from the design of the array, the majority of the probe CpG targets were located in genic regions. While *cis* mQTL methylation probes showed no large deviation in genomic annotation from all probes, the number of *trans* mQTL CpGs was substantially inflated in both upstream and 5’ UTR regions.

**Table 1:** Genomic annotation of mQTL CpG site locations. Only categories from ANNOVAR that contain greater than 1% of probes are included. A substantial inflation of “Upstream” and “UTR5” is found for probes with *trans* mQTL.

| Classification | All Array Probes | *Cis* mQTL Probes | *Trans* mQTL probes |
| --- | --- | --- | --- |
| Intronic | 33.7% | 36.0% | 28.1% |
| Intergenic | 21.3% | 25.8% | 14.5% |
| Upstream | 19.2% | 17.2% | **29.5%** |
| Exonic | 9.0% | 6.6% | 7.1% |
| UTR5 | 6.0% | 3.4% | **13.2%** |
| UTR3 | 3.8% | 3.6% | 1.7% |
| ncRNA-intronic | 2.5% | 2.9% | 1.4% |
| ncRNA-exonic | 1.5% | 1.4% | 1.8% |

### Role of mQTL in Complex Traits and Disease

To assess the role of mQTL in driving the phenotypic variation of complex traits and disease, we used LD Score regression ^26,27^ to partition the trait heritability into components due to mQTL and the rest of the genome. LD Score regression uses summary statistics from GWAS, allowing us to investigate a range of traits and diseases using results from large consortia (for height ^28^, BMI ^29^, schizophrenia ^30^, ulcerative colitis ^31^, Crohn’s disease ^31^, coronary artery disease ^32^, type 2 diabetes ^33^, rheumatoid arthritis ^34^, and educational attainment ^35^).

The replicated mQTL were firstly filtered to have no SNP pairs with an estimated r^2^ of greater than 0.8. This allows for straightforward generation of sets of SNPs to estimate the distribution of variance explained under the null hypothesis, as then the LD structure is similar to that of a random set of minor allele frequency matched SNPs. Two different null hypotheses were used. The first (null #1) accounted for the fact that on average SNPs with a higher heterozygosity explain more variation in a trait by drawing random sets of SNPs with a matched minor allele frequency (in bins of 0.05 width). The second (null #2) in addition matched the genomic location of randomly sampled SNPs using annotation from ANNOVAR ^36^. This accounts for the observation that a large proportion of the genetic variation in complex traits is explained by genic regions and that the array (and thus *cis* mQTL locations) is very gene centric.

Under null #1, height, ulcerative colitis, Crohn’s disease and coronary artery disease all showed a significant inflation of the proportion of genetic variation explained by mQTL (Table 2), although none of these were significant after accounting for the genomic location of the mQTL SNP (null #2). However, sets of SNPs generated for null #2 tag many of the same regions of the genome as the mQTL SNP due to large number of genic mQTL identified in this study compared to genes in the genome. Thus it is not surprising that none of the tests under null #2 are significant, and we cannot distinguish between the hypotheses of close linkage and causality. It is of note that all of those tests that were significant under null #1 explained more than average variation under null #2.

**Table 2:** LDScore regression partitioning of the heritability for a variety of traits and disease. For each trait, the heritability was partitioned into components explained by mQTL and the rest of the genome and the proportion of the total explained heritability attributable to mQTL was calculated. Several phenotypes showed a significant role of mQTL under the first null hypothesis (matched allele frequencies) but these did not remain significant when SNPs were matched to genomic location (Null #2).

| Trait | SNP | N * | mQTL Proportion | Null #1 |  | Null #2 |  |
| --- | --- | --- | --- | --- | --- | --- | --- |
|  |  |  |  | Mean (S.E.) | P-value | Mean (S.E) | P-value |
| Height | 2,517,431 | 253,288 | 0.330 | 0.083 (0.040) | $2.1 \times 10^{-10}$ | 0.269 (0.052) | 0.12 |
| BMI | 2,524,366 | 322,154 | 0.245 | 0.206 (0.084) | 0.32 | 0.303 (0.096) | 0.73 |
| Schizophrenia | 6,101,975 | 82,315<sup>†</sup> | 0.262 | 0.152 (0.046) | 0.0098 | 0.271 (0.047) | 0.57 |
| Ulcerative colitis** | 1,346,293 | 27,432 | 0.333 | 0.071 (0.064) | $2 \times 10^{-5}$ | 0.299 (0.094) | 0.37 |
| Crohn's Disease** | 948,687 | 20,883 | 0.305 | 0.053 (0.048) | $6 \times 10^{-8}$ | 0.252 (0.071) | 0.23 |
| Coronary Artery Disease | 2,398,186 | 86,995 | 0.292 | 0.038 (0.058) | $5.5 \times 10^{-6}$ | 0.238 (0.076) | 0.24 |
| Type 2 Diabetes | 2,411,307 | 80,788 | 0.297 | 0.172 (0.106) | 0.12 | 0.253 (0.095) | 0.32 |
| Rheumatoid Arthritis** | 8,409,120 | 58,284 | 0.136 | 0.087 (0.104) | 0.32 | 0.202 (0.127) | 0.70 |
| Educational Attainment | 2,291,668 | 126,559 | 0.110 | 0.114 (0.062) | 0.52 | 0.227 (0.073) | 0.94 |

\* N = N\_cases + N\_controls for case-control studies.
\*\* Excluding the HLA region of chromosome 6
<sup>†</sup> Contains non-European samples

Due to the limitations of the genomic partitioning, a second approach to investigate the effect of mQTL on complex traits and disease was taken. If mQTL are a driving force behind phenotypic variation, then it would be expected that mQTL SNPs with large effects on DNA methylation would also have large effects on the complex trait. To test this, we estimated the correlation between the mQTL SNP effect size and its effect from the large GWAS studies. The absolute value of the effect (or log odds-ratio) on both DNA methylation and the trait was used as it is expected that there will be variation in whether DNA methylation is protective or not for different regions of the genome. In addition, the effect sizes were corrected for the expected relationship between effect size and minor allele frequency by multiplying the effect size by, where *f* is the minor allele frequency of the SNP. After correcting for minor allele frequency, no significant correlation was observed between the effects sizes of the SNPs on the mQTL and the corresponding SNP effect sizes on any of the tested traits (Table S2).

## DISCUSSION

We have identified 52,916 *cis* and 2,025 *trans* mQTL that are replicated across two independent cohorts at very stringent significance levels. While the mQTL can explain a large proportion of the genetic variation underlying DNA methylation variation, there is still substantial genetic variation remaining to be explained. Using the twin family structure in the Brisbane Systems Genetics Study, we have previously shown that the average heritability of DNA methylation at sites measured by the Illumina HumanMethylation450 array is 0.187 ^6^. The average proportion of phenotypic variation explained by all mQTL across all DNA methylation probes in this study (including probes that had no mQTL and thus explained zero variation) is 0.021. Thus, the mQTL identified here explain approximately 11.2% of the total genetic variation for DNA methylation. This implies there is substantial genetic variation for DNA methylation remaining to be discovered through additional variants in *cis* and/or many more *trans* variants with small effects in larger samples.

By partitioning heritability into components due to mQTL SNPs and the rest of the genome, we established that the identified mQTL explained a significant amount of the genetic variation for a number of complex traits and diseases. Using a null distribution generated by randomly sampling SNPs from the genome with matching minor allele frequencies showed significant amounts of genetic variation were explained by mQTL for height, schizophrenia, ulcerative colitis, Crohn’s disease, and coronary artery disease. This enrichment of mQTL in disease associated regions was explained by the genomic location of the mQTL SNP. This is due to most mQTL SNP being *cis* to the DNA methylation probes, which also tend to be found in genic regions due to the design of the array, combined with the observation that genic regions explain more of the heritability for many traits ^21^. Previous studies that have shown a relationship between mQTL and bipolar disorder ^11^ and schizophrenia ^18^ QTL whilst only considered MAF when sampling SNPs for the null distribution, and, as demonstrated here, the results are likely to be driven by the common genomic function of the SNPs. Testing for a role of mQTL in complex traits and disease beyond that explained by genomic location is difficult due to the large number of mQTL replicated in this study. This means that a large proportion of genes in the genome are tagged by an mQTL and any null sample of SNPs will cover many of the same genomic regions. This makes any test for the proportion of heritability explained by mQTL being extremely conservative.

Determining whether associations detected in the same genetic region for DNA methylation and a disease are the result of (mediated) pleiotropy or just close linkage is a difficult prospect. To have potential for pleiotropy, the set of potential causal variants for the two associations will need to overlap. Fine-mapping to a set of potential causal variants can be determined by statistical prioritisation using only association statistics ^37–39^, or in combination with other genomic data ^40–42^. Reducing the set of potential causal variant(s) underlying a mQTL using these approaches is helped by the large amount of phenotypic variation the mQTLs explain. There is also strong potential to determine causal SNPs for mQTLs in cell lines using CRISPR genome editing ^43^ as the end phenotype is directly observable in the cell, unlike the case for complex traits and disease where a phenotype to investigate in cell lines is generally unclear.

We observed a strong over-representation of *trans* mQTL SNP in the 1Mbp subtelomeric region of the genome, as had been previously noted ^17^. No association of the *trans* mQTL SNP or methylation probes was found with telomere length in the LBC1936 cohort. The *trans* mQTLs were significantly inflated for methylation probes found in the upstream regions of genes, indicating a potential effect on the regulation of gene-expression. However, there was no overlap with *trans* eQTLs identified in the BSGS ^22^. The mechanism and potential importance of subtelomeric regions in altering DNA methylation throughout the genome warrants further investigation and at this stage artefacts of the technology cannot be excluded.

In summary, we have identified and replicated a large number of genetic loci associated with DNA methylation in both *cis* and *trans*. We demonstrated an overlap of mQTL and loci for complex traits and diseases, which was explained by the genomic location of the mQTL SNPs.

## MATERIALS AND METHODS

### Brisbane Systems Genetics Study (BSGS)

DNA methylation was measured on 614 individuals from 177 families of European descent recruited as part of a study on adolescent twins and selected from individuals in the Brisbane Systems Genetics Study ^6,22^. Families consist of adolescent monozygotic (MZ) and dizygotic (DZ) twins, their siblings, and their parents. DNA was extracted from peripheral blood lymphocytes by the salt precipitation method ^44^. The BSGS study was approved by the Queensland Institute for Medical Research Human Research Ethics Committee. All participants gave informed written consent.

### Lothian Birth Cohorts

Methylation data were analysed from the combined data of the Lothian Birth Cohort 1921 (LBC1921) and the Lothian Birth Cohort 1936 (LBC1936) ^23–25^. The LBC1921 and LBC1936 are longitudinal studies of ageing, with a focus on cognition, in groups of initially healthy older people. DNA methylation was measured in 446 LBC1921 subjects at an average age of 79 years, and in 920 LBC1936 subjects at an average age of 70 years ^45^. Following informed consent, venesected whole blood was collected for DNA extraction by standard methods in both LBC1921 and LBC1936. Ethics permission for the LBC1921 was obtained from the Lothian Research Ethics Committee (Wave 1: LREC/1998/4/183). Ethics permission for the LBC1936 was obtained from the Multi-Centre Research Ethics Committee for Scotland (Wave 1: MREC/01/0/56), the Lothian Research Ethics Committee (Wave 1: LREC/2003/2/29). Written informed consent was obtained from all subjects.

### DNA Methylation

DNA methylation was measured using Illumina HumanMethylation450 BeadChips as described in detail elsewhere ^6,45^. The HM 450 BeadChip-assessed methylation status was interrogated at 485,577 CpG sites across the genome. It provides coverage of 99% of RefSeq genes. Methylation scores for each CpG site are obtained as a ratio of the intensities of fluorescent signals and are represented as β-values. DNA methylation data for the BSGS is available at the Gene Expression Omnibus under accession code GSE56105, and the LBC data is available at the European Genome-phenome Archive under accession number EGAS00001000910.

Probes on the sex chromosomes or having been annotated as binding to multiple chromosomes ^46^ were removed from the analysis, as were non CpG sites. Probes with excess missingness or high numbers of individuals with detection p-value less than 0.001 were also removed. After cleaning, 397,710 probes remained for association analysis in both cohorts.

### Normalisation

Array data were background corrected, followed by individual probes being normalised using a generalised linear model with a logistic link function. Corrections were made for the effects of chip (which encompasses batch processing effects), position on the chip, sex, age, age^2^, sex x age and sex x age^2^. In addition, the LBC data were corrected for white blood cell counts (basophils, eosinophils, monocytes, lymphocytes, and neutrophils). The LBC data were normalised for the two cohorts individually before combining the data for further analysis.

Outlying data points can result in a high number of false positive in GWAS analysis when associated with rare variants. To address this, the BSGS cohort removed any measurement at a probe that was greater than five interquartile ranges from its nearest quartile. In the LBC, probes that had such outliers were restricted to testing association with SNPs having a minor allele frequency greater than 5%.

### Genotyping and Imputation

Both the BSGS and LBC were genotyped on Illumina 610-Quad Beadchip arrays, with full details of genotyping procedures described elsewhere ^47,48^. After standard quality control, the BSGS and LBC had 528,509 and 549,692 SNPs remaining respectively.

The remaining genotyped SNPs were phased using SHAPEIT ^49,50^ and imputed against 1000 Genomes Phase I Version 3 ^51,52^ using Impute V2 ^53,54^. Raw imputed SNPs were filtered to remove any SNPs with low imputation quality as defined by an r^2^ < 0.8. Subsequent quality control removed SNPs with MAF < 0.05, and those with HWE p < 1 x 10^−6^. The “best-guess” (highest probability) genotype was used for the GWAS analyses.

### Genome-Wide Association Analysis

Genome-wide association (GWAS) was performed individually on the BSGS and LBC cohorts, with each serving as an independent discovery cohort and replication performed in the other.

To reduce the massive computational burden, GWAS was performed in two stages. Firstly the *cis* region to the methylation probe – defined as a window 2Mbp each side of the target CpG site location – was investigated. A significance threshold of 10^−11^ was used, which is a stringent p=0.05 Bonferroni correction for the approximate number of independent SNPs in the window and number of probes analysed. Significant associations were replicated with a Bonferonni corrected (based on the approximate number of independent mQTL) p-value of 10^−6^ and having effect in the same direction in the other sample. When a single methylation probe had a replicated association from both cohorts but at a different SNP, the SNP with the best combined evidence of association was selected for further analyses.

Association with *trans* SNPs (defined as all SNPs outside the 4Mbp window used in the *cis* analysis) was performed in two steps. Firstly, all chromosome/probe pairs were analysed on non-imputed genotyped data, which reduced the number of tests performed by a factor of 10. This was particularly important for the BSGS cohort which had related individuals and thus was much slower to analyse. Any chromosome/probe pair that had an association at p<10^−7^ was then reanalysed using imputed SNP data. An experiment-wide significance of 10^−13^ was used for *trans* associations, which is the standard GWAS genome-wide significance threshold of 5 x 10^−8^ Bonferroni corrected for the number of probes tested. The replication threshold of 10^−5^ was used, again being more stringent than a 5% significance Bonferroni corrected for the number of associations to be replicated.

Association testing was performed using MERLIN ^55^ using the –fastAssoc option for the BSGS cohort (to account for family structure) and PLINK ^56^ for the combined LBC cohorts.

### Genomic Annotation of SNP and Methylation Sites

SNPs and the CpG targets of methylation probes were functionally annotated using ANNOVAR ^36^, using the hg19 annotation with the distance of the upstream and downstream regions of genes being 2Mbp to align with our definition of *cis* loci.

### Telomere Measurements

Telomere length was measured using the same blood sample as methylation in the LBC1936 cohort using a quantitative real-time polymerase chain reaction (PCR) assay ^57^. The intra-assay coefficient of variation was 2.7% and the inter-assay coefficient of variation was 5.1%. Four internal control DNA samples were run within each plate to correct for plate-to-plate variation. These internal controls are cell lines of known absolute telomere length whose relative ratio values (telomere starting quantity/glyceraldehyde 3-phosphate dehydrogenase starting quantity) were used to generate a regression line by which values of relative telomere length for the actual samples were converted into absolute telomere lengths. Measurements were performed in quadruplicate and the mean of the measurements used. PCRs were performed on an Applied Biosystems (Pleasonton, CA, USA) 7900HT Fast Real Time PCR machine.

### Partitioning Heritability

The heritability of a trait explained by all GWASed SNPs was partitioned in to a component due to all discovered mQTL and all remaining SNP using LD Score regression ^26,27^. The sum of the LD *r*^2^ values for between that target SNP and all other SNPs within the 1Mbp region centred on the target SNP ^58^, and was calculated using the European samples from the 1000 Genomes project ^51,52^ using the software GCTA (–ld-score option) ^59^. The LD score at a SNP, *j*, is then calculated as:

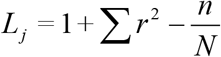

where *n* is the number of SNP in the window and *N* is sample size used to calculate the *r*^2^ measures.

Using the summary statistics from a large GWAS for a quantitative trait or disease, the heritability of the trait is partitioned into components due to mQTL and the rest of the genome using a regression

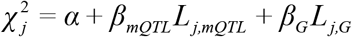

where is the chi-square test statistics for SNP *j*. The heritability attributable to mQTL is calculated as

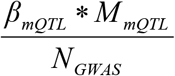

where *M_mQTL_* is the number of mQTL SNPs and *N_GWAS_* is the sample size of the GWAS from which the summary statistics were obtained. The heritability attributable to the rest of the genome is calculated similarly.

## ACKNOWLEDGEMENTS

We thank the cohort participants and team members who contributed to these studies. The Brisbane Systems Genetics Study (BSGS) was supported by NHMRC grants 1010374, 496667, 1046880.A.F.M., J.E.P., N.R.W., P.M.V., and G.W.M. are supported by the NHMRC Fellowship Scheme (1083656, 1107599, 1078901, 1078037 and 1078399) and grants (1050218). J.Y. is supported by the Sylvia & Charles Viertel Charitable Foundation. We acknowledge funding by the Australian Research Council (A7960034, A79906588, A79801419, DP0212016, DP0343921), and the Australian National Health and Medical Research Council (NHMRC) Medical Bioinformatics Genomics Proteomics Program (grant 389891) for building and maintaining the adolescent twin family resource through which samples were collected. Phenotype collection in the Lothian Birth Cohort 1921 (LBC1921) was supported by the UK’s Biotechnology and Biological Sciences Research Council (BBSRC), The Royal Society and The Chief Scientist Office of the Scottish Government. Phenotype collection in the Lothian Birth Cohort 1936 (LBC1936) was supported by Age UK (The Disconnected Mind project). Genotyping of LBC1921 and LBC1936 was funded by the BBSRC. Methylation typing of LBC1921 and LBC1936 was supported by The Centre for Cognitive Ageing and Cognitive Epidemiology (Pilot Fund award), Age UK, The Wellcome Trust Institutional Strategic Support Fund, The University of Edinburgh, and The University of Queensland. Telomere length data was generated with the support of Carmen Martin-Ruiz and Thomas von Zglinicki. REM, SEH, JMS, IJD and PMV are members of the University of Edinburgh Centre for Cognitive Ageing and Cognitive Epidemiology (CCACE). CCACE is supported by funding from the BBSRC, the Economic and Social Research Council (ESRC), the Medical Research Council (MRC), and the University of Edinburgh as part of the cross-council Lifelong Health and Wellbeing initiative (MR/K026992/1).

## AUTHOR CONTRIBUTIONS STATEMENT

Conceived and designed the experiments: AFM, REM, NRW, IJD, PMV, GWM. Performed the experiments: SEH, JG, AKH, LB, JNP, LM. Analyzed the data: AFM, REM, SS, JY. Contributed reagents/materials/analysis tools: SEH, JG, AKH, NGM, JMS, LM Wrote the paper: AFM, REM, NRW, IJD, PMV, GWM. All authors read and approved the final manuscript.

## ADDITIONAL INFORMATION

### Data Availability

DNA methylation data for the BSGS is available at the Gene Expression Omnibus under accession code GSE56105, and the LBC data is available at the European Genome-phenome Archive under accession number EGAS00001000910.

